# Statistical learning of distractor co-occurrences facilitates visual search

**DOI:** 10.1101/2022.04.20.488921

**Authors:** Sushrut Thorat, Genevieve Quek, Marius V. Peelen

## Abstract

Visual search is facilitated by knowledge of the relationship between the target and the distractors, including both where the target is likely to be amongst the distractors and how it differs from the distractors. Whether the statistical structure amongst distractors themselves, unrelated to target properties, facilitates search is less well understood. Here, we assessed the benefit of distractor structure using novel shapes whose relationship to each other was learned implicitly during visual search. Participants searched for target items in arrays of shapes that comprised either four pairs of co-occurring distractor shapes (structured scenes) or eight distractor shapes randomly partitioned into four pairs on each trial (unstructured scenes). Across five online experiments (N=1140), we found that after a period of search training, participants were more efficient when searching for targets in structured than unstructured scenes. This structure-benefit emerged independently of whether the position of the shapes within each pair was fixed or variable, and despite participants having no explicit knowledge of the structured pairs they had seen. These results show that implicitly learned co-occurrence statistics between distractor shapes increases search efficiency. Increased efficiency in the rejection of regularly co-occurring distractors may contribute to the efficiency of visual search in natural scenes, where such regularities are abundant.

## Introduction

Visual search is the task of finding a target object (e.g., a computer mouse on a desk) amongst distractor objects (e.g., other objects on the desk). It is well-established that search-difficulty (as measured by reaction time and/or accuracy) increases linearly with the number of distractors (Wolfe, 1998). Interestingly, this relationship is much weaker for search in natural scenes than for search in artificial arrays comprising randomly arranged objects (Wolfe, Alvarez, et al., 2011). What makes naturalistic visual search so efficient?

An important contribution comes from the information scene context provides about spatial (“where”) and featural (“what”) target properties (Castelhano & Krzyś, 2020; Oliva & Torralba, 2007; Peelen & Kastner, 2014; Võ et al., 2019; Wolfe, Võ, et al., 2011). For example, the likely location of the target in a scene can be learned and used to facilitate search, both based on recent experience in controlled laboratory experiments (“contextual cueing”; Chun, 2000) and based on long-term daily-life experience (Castelhano & Krzyś, 2020; Võ et al., 2019): when searching for a computer mouse, we start searching to the right of the keyboard and below the monitor. Scene context also provides information about the features that characterize the target (Peelen & Kastner, 2014), or distinguish the target from the distractors (Geng & Witkowski, 2019): we look for a small target far away and a large target nearby (Gayet & Peelen, 2022). Finally, targets are recognized more quickly when embedded in context, reflecting the facilitatory influence of contextual expectations on object recognition (Bar, 2004; de Lange et al., 2018). Thus, our long- and short-term experience with regularities in where and how targets appear in scenes contributes to the efficiency of visual search.

Importantly, real-world scenes are additionally characterized by regularities amongst distractors themselves. For example, when searching for the television in a living room, the co-occurrence statistics and spatial arrangements of many distractor objects (e.g., chairs, tables, lamps) is relatively stable. Some of these regularities are consistent across environments and learned across a lifetime (e.g., a typical living room layout), while others are specific to a particular context and learned more rapidly (e.g., the arrangement of objects in my friend’s living room). Previous research found that visual search is easier when distractor objects are arranged in configurations that follow real-world regularities (e.g., lamp above table) than when they are arranged in unfamiliar configurations (e.g., lamp below table; Kaiser et al., 2014). These results may reflect more efficient encoding of familiar object pairs based on long-term experience (Bar & Ullman, 1996; Quek & Peelen, 2020; Stein et al., 2015), facilitating visual search when these objects appear as distractors. Alternatively, visual search may be disrupted when distractor configurations violate higher-level functional and semantic associations (Spaak et al., 2022; Võ & Wolfe, 2013).

Together, the findings reviewed above raise the question of whether statistical regularities amongst distractors contribute to the efficiency of search, independently of target-distractor regularities and independently of long-term semantic knowledge. Interestingly, previous research has shown that statistical regularities between shapes can be learned rapidly (Fiser & Aslin, 2001; Fiser & Lengyel, 2019; Schapiro & Turk-Browne, 2015). For example, when participants passively view displays in which one shape frequently appears together with another shape (always in the same configuration), participants later report higher familiarity for these pairs relative to control pairs (Fiser & Aslin, 2001). Furthermore, such newly-learned shape pairs show object-like behavioral signatures, with attention spreading from one shape to the other (Lengyel et al., 2021), akin to effects of perceptual grouping (Egly et al., 1994; Scholl, 2001). If such regularity-based object grouping occurs amongst distractor objects during visual search, this compression could effectively reduce the distractor numerosity (Zhao & Yu, 2016), thereby enhancing search performance, similar to how perceptual grouping facilitates visual search (Donnelly et al., 1991; Humphreys et al., 1989; Rauschenberger & Yantis, 2006).

To test whether newly-learned statistical regularities amongst distractors contribute to the efficiency of search, here we combined a statistical learning paradigm with a visual search task using novel shapes. The use of novel shapes allowed for full control over co-occurrence probabilities and low-level stimulus properties. Participants searched for pre-cued target shapes in arrays that consisted of either four pairs of co-occurring distractor shapes (structured scenes) or else eight distractor shapes randomly partitioned into four pairs (unstructured scenes). Participants were not informed about the co-occurrences, such that all co-occurrence statistics were learned during the search task itself. To assess if the specific spatial arrangement of co-occurring shapes within the pairs was essential for distractor complexity reduction, the co-occurring shapes either had fixed arrangements (e.g., shape A always appeared above shape B) or their locations within the pairs could be swapped (e.g., shape A could appear above or below shape B).

Across multiple experiments, we found that participants were more efficient in searching for targets in the structured scenes than the unstructured scenes. Interestingly, this pattern was independent of whether the arrangement of co-occurring shapes within the pairs was fixed or not. Finally, unlike previous statistical learning studies where the co-occurring objects were attended (Fiser & Aslin, 2001), here participants were not able to indicate which shapes co-occurred during the visual search experiment, indicating that statistical regularities in the environment facilitate search even when these regularities are not explicitly noticed.

## Materials and Methods

### Participants

Participants were recruited online using Prolific, received monetary compensation for their participation, and provided informed consent before starting the experiment. The study was approved by the Radboud University Faculty of Social Sciences Ethics Committee (ECSW2017–2306-517) and was carried out in accordance with the provisions of the World Medical Association Declaration of Helsinki. Participants from whom we obtained partial data were excluded from the analysis (∼10% dropout rate). For any given experiment requiring a particular number of participants (see below), we first tested around that number of participants balancing the blocking order of scene structure. Then, participants whose overall accuracy and reaction times were above or below 3 standard deviations (SDs) from the means were removed. This was done iteratively until no exclusions happened. Then, more participants were added to get to the desired number and this exclusion process was repeated. In the end, we obtained the desired number of participants for each experiment whose accuracies and reaction times (for correct responses) were within 3 SDs from the means and the blocking order was balanced.

The desired number of participants for the two initial experiments was 40: Experiment 1A (mean age: 25.3 years, SD = 4.4), Experiment 1B (mean age: 27.1 years, SD = 4.7). The number of participants for the two large-scale replication experiments was 400: Experiment 2A (mean age: 24.1 years, SD = 4.3), Experiment 2B (mean age: 25.6 years, SD = 6.4). Finally, the number of participants for Experiment 3 was 260 (N=260; mean age: 23.9 years, SD = 4.5). This experiment was pre-registered (https://aspredicted.org/blind.php?x=5ne7qa).

### Stimuli

The stimulus set consisted of 20 novel shapes (See Figure 1 for examples), a subset of which overlapped with those from seminal statistical learning studies (Fiser & Aslin, 2001, 2005). For each participant, we randomly assigned the 20 shapes to three different sets that were maintained throughout the experiment: 4 shapes were used as search targets (target set), 8 shapes were allocated into 4 co-occurring distractor pairs (structured distractor set), and 8 shapes were used to create 4 random distractor pairs on each new trial (unstructured distractor set). Critically, a shape assigned to the structured set only ever appeared in a vertical pairing with its nominated partner shape. In *fixed* arrangements (Experiments 1A, 2A, 3), the shapes in the structured set appeared in specific vertical arrangements throughout the experiment (e.g., shape A always appeared above shape B). In *free* arrangements (Experiments 1B, 2B), the shapes in the structured set randomly appeared in one of two vertical arrangements across trials (e.g., shape A could appear either above or below shape B). In contrast to these structured conditions, shapes assigned to the unstructured set could be paired with any other shape from the unstructured set and could occupy either the top or bottom position within this random pairing.

**Figure 1:**
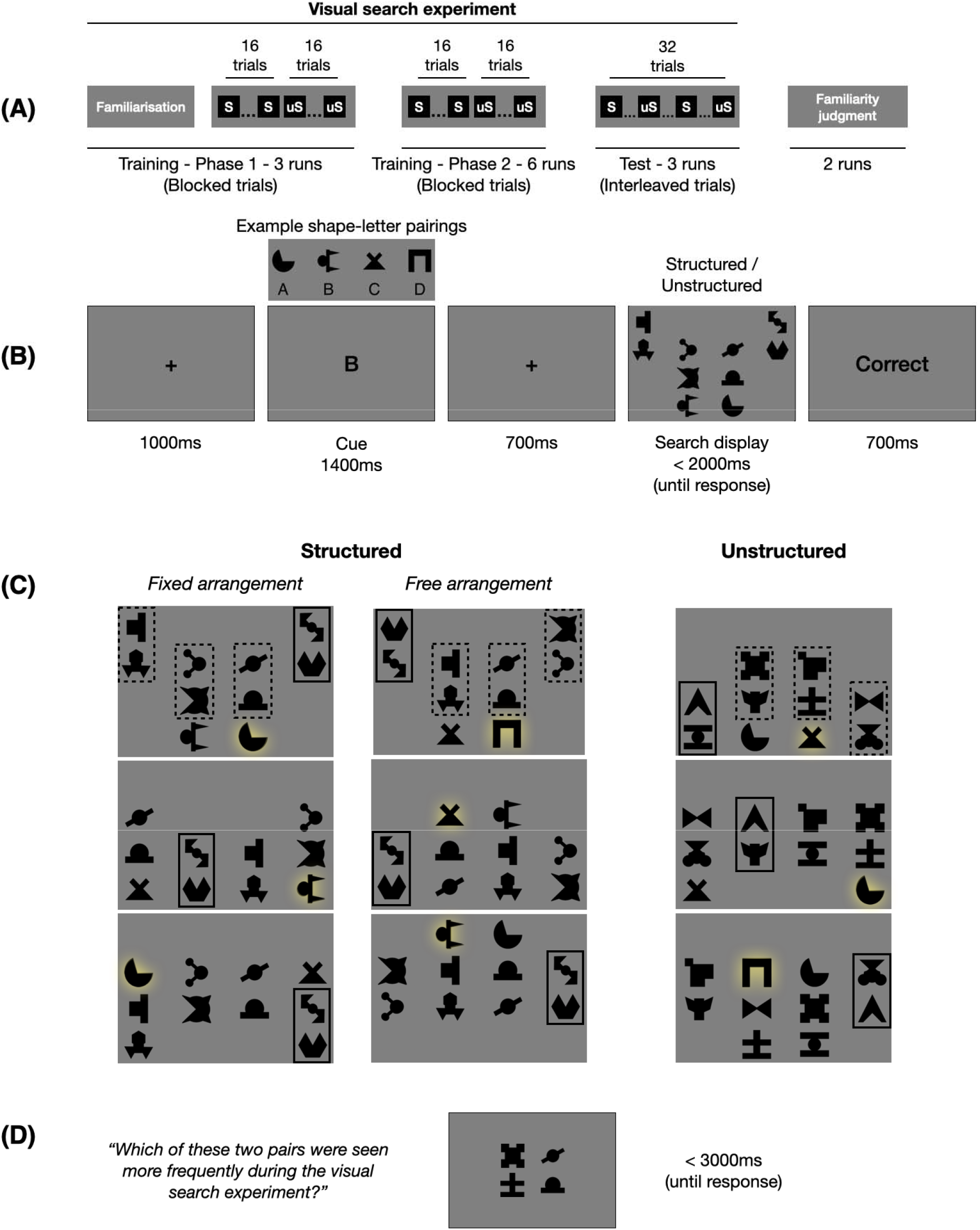
Experimental procedure and design. (A) Schematic outline of the experiment. S = structured scenes; uS = unstructured scenes. Structured and unstructured scenes were blocked during the first nine training runs of the visual search experiment but interleaved during the final three test runs. The visual search experiment was followed by a familiarity judgment task. (B) The trial structure of the visual search experiment. Participants had to search for a target shape cued by its corresponding letter in the subsequent search display and indicate if the target was present on the left or the right part of the display within 2s. (C) Example layouts for the structured and unstructured visual search displays for one participant. Ten shapes appeared on each trial: 8 distractors, 1 target (highlighted in yellow, color not shown during the experiment), and 1 foil (which could be a target on other trials). The distractors were presented as 4 pairs (indicated by the dashed outlines in the first example; one pair is outlined across displays to illustrate the respective manipulation (fixed, free) across trials). In the structured scenes, the distractors co-occurred in pairs of two (with either fixed arrangements within the pairs or not, in separate experiments). In the unstructured scenes, the distractors were randomly partitioned into four pairs on each trial. Search performance was compared between structured and unstructured scenes. (D) An example trial of the familiarity judgment task. Participants had to judge which of the two vertical pairs (one taken from the structured scenes and the other from the unstructured scenes) had been seen more frequently during the visual search experiment.

The search display was 16em x 28em, where em is the font size on the participant’s display. This size was chosen such that the display would approximately extend around 6 degrees of visual angle during typical viewing conditions. We reasoned that those participants who used smaller screens also had smaller font sizes and were positioned closer to the screen, such that the visual angle subtended by the relevant stimuli were roughly equated across screen sizes. Note, however, that because the study was conducted online, we could not fully control the visual angle subtended by the search display. The experiment was programmed in JavaScript with jsPsych and hosted online on Pavlovia (Open Science Tools Limited, 2021).

### Procedure and design

Each trial of the visual search task started with a letter cue (1400 ms) indicating the target shape for that trial (Figure 1B). After a brief (700 ms) delay, a search display with 10 shapes appeared. Participants used the keyboard to indicate whether the target was present on the left (“F” key) or the right (“J” key) side of the display.

Each search display consisted of 4 distractor shape pairs, the target shape, and a foil shape (one of the other three target shapes not currently being searched for) arrayed symmetrically on a 4×4 grid (Figure 1C). The distractor pairs could comprise either the 4 distractor pairs from the structured set or else 4 randomly generated distractor pairs from the unstructured set. Thus, on each trial, participants searched for the target in either a structured or unstructured scene. One distractor pair appeared in each of the 4 columns of the grid, in random horizontal order. The vertical position of the pairs was random, but with the constraint that the locations were mirrored horizontally. The target appeared in one of the remaining locations vertically adjacent to a pair, with the foil (one of the other 3 targets) in the horizontally-mirrored location. The location randomization process ensured that the probability of the target’s location was uniform across the entire grid.

Participants completed a total of 12 runs of the visual search task (Figure 1A). Each run consisted of 32 trials (16 trials with structured distractor pairs and 16 trials with unstructured distractor pairs), for a total of 384 trials. Structured and unstructured trials were blocked in the first 9 runs (training runs), but randomly interleaved in the last 3 runs (test runs; Figure 1A). The order of blocking (structured trials first or unstructured trials first) was maintained for a participant throughout the experiment and balanced across participants. We elected to block the structure conditions during training based on evidence that humans appear to learn statistical associations faster when these are presented in a blocked rather than interleaved order (Flesch et al., 2018). All analyses focused on responses in the interleaved test runs to avoid possible block-based differences in arousal or strategy.

The experiment started with three training runs to familiarize participants with the target letter-shape association and to practice the visual search task (phase 1 training; Figure 1A). We used letter cues (rather than target picture cues) to increase the difficulty of the task: target picture cues would perceptually prime the target, reducing the influence of distractors on search performance (Wolfe et al., 2004; Schmidt & Zelinsky, 2009). These runs started with a familiarization block where the letters and their associated target shapes were shown sequentially four times. Next, participants completed six runs of the visual search task to (implicitly) learn the statistical regularities of the structured distractor pairs (phase 2 training; Figure 1A). These runs started with instructions and a reminder of the target letter-shape associations to refresh participants’ memory. Finally, participants completed 3 test runs where the structured and unstructured conditions were interleaved.

### Familiarity judgment task

After completing the 12 runs of the visual search task, participants additionally completed two runs of a two-alternative-forced-choice (2AFC) familiarity judgment task (Fiser & Aslin, 2001). This component of the experiment aimed to assess participants’ explicit knowledge about the shape pairs that had appeared as distractors during the main visual search experiment. On each trial, participants were asked to indicate (or guess) within three seconds which of two shape-pairs had appeared more frequently during the visual search experiment (Figure 1D). One of the pairs was a structured pair, taken directly from the preceding visual search component, while the other was an unstructured pair. We compared the 4 structured pairs with 4 randomly selected unstructured pairs, which were held constant throughout the familiarity judgment task, such that within the familiarity judgment task all pairs were presented equally often. The position of the shapes within the pairs was also held constant during the familiarity judgment task. Importantly, in the visual search task, the structured pairs had been presented 14 times more often than the unstructured pairs; if participants noticed these regularities, they should show above-chance performance on the familiarity judgment task (Fiser & Aslin, 2001).

Each run contained trials showing the 16 possible combinations between the 4 original structured pairs and the 4 selected unstructured pairs. The main analyses focused on these trials, as they provide the most sensitive test of familiarity and were included in all familiarity judgment experiments. The familiarity judgment experiments for the fixed arrangement condition (Experiments 2A and 3) additionally included trials in which either the position or the partner was swapped across the set of structured pairs (and, separately, the set of selected unstructured pairs). Four partner-swapped pairs were constructed, separately for the structured and the unstructured scenes, by swapping the partners of the shapes while maintaining their relative positions in the pairs. Four position-swapped pairs were constructed, separately for the structured and the unstructured scenes, by swapping the positions of the shapes within their pairs. These two manipulations led to 32 additional comparisons between the shapes from the structured and unstructured scenes, for a total of 48 trials per run.

For the first 200 participants in Experiment 2A, these trials were presented in random order and feedback was provided at the end of each run. For the last 200 participants of Experiment 2A and for all participants in Experiment 3, the original 16 comparisons were presented at the beginning of the run, with the other conditions presented interleaved in the remainder of the run. For these participants, feedback was only provided at the end of the second run. Only participants were included who responded at least once to each condition in each run, leaving 368 of the 400 participants in Experiment 2A for the familiarity judgment analysis. Participants not meeting this requirement in Experiment 3 were replaced, such that all 260 participants were included in the familiarity judgment analysis. In Experiment 2B (free arrangement), the first half of the participants (N = 200) did not complete the familiarity judgment task. For the second half of the participants (N = 200), in each run, only the 16 original comparisons between the 4 forced pairs from the structured scenes and 4 forced pairs from the unstructured scenes were shown. All participants responded at least once in each run.

### Data availability

The analysis code and data accompanying these experiments can be found on OSF, https://doi.org/10.17605/OSF.IO/EM2XF

## Results

### Search performance as a function of learned distractor structure

To test whether the presence of co-occurring distractors facilitated visual search, we evaluated the difference between search performance in structured scenes and unstructured scenes in terms of both accuracy and reaction time in the test runs (i.e., after a period of exposure during training). Reaction time was computed for correct trials only. Trials with reaction times below 300ms or above 2000ms were not included in the analyses. The difference in performance between structured and unstructured scenes was termed the *structure-benefit* (indicated by a higher search accuracy or faster reaction times in the structured scenes). In addition to accuracy and reaction time, we used the inverse efficiency score (IES = average reaction time / average accuracy) as a combined measure for the structure-benefit. IES is a useful measure when accuracy is high (>90%) and effects in accuracy and reaction time go in the same direction (Bruyer & Brysbaert, 2011), as was the case here (Figure 2).

**Figure 2:**
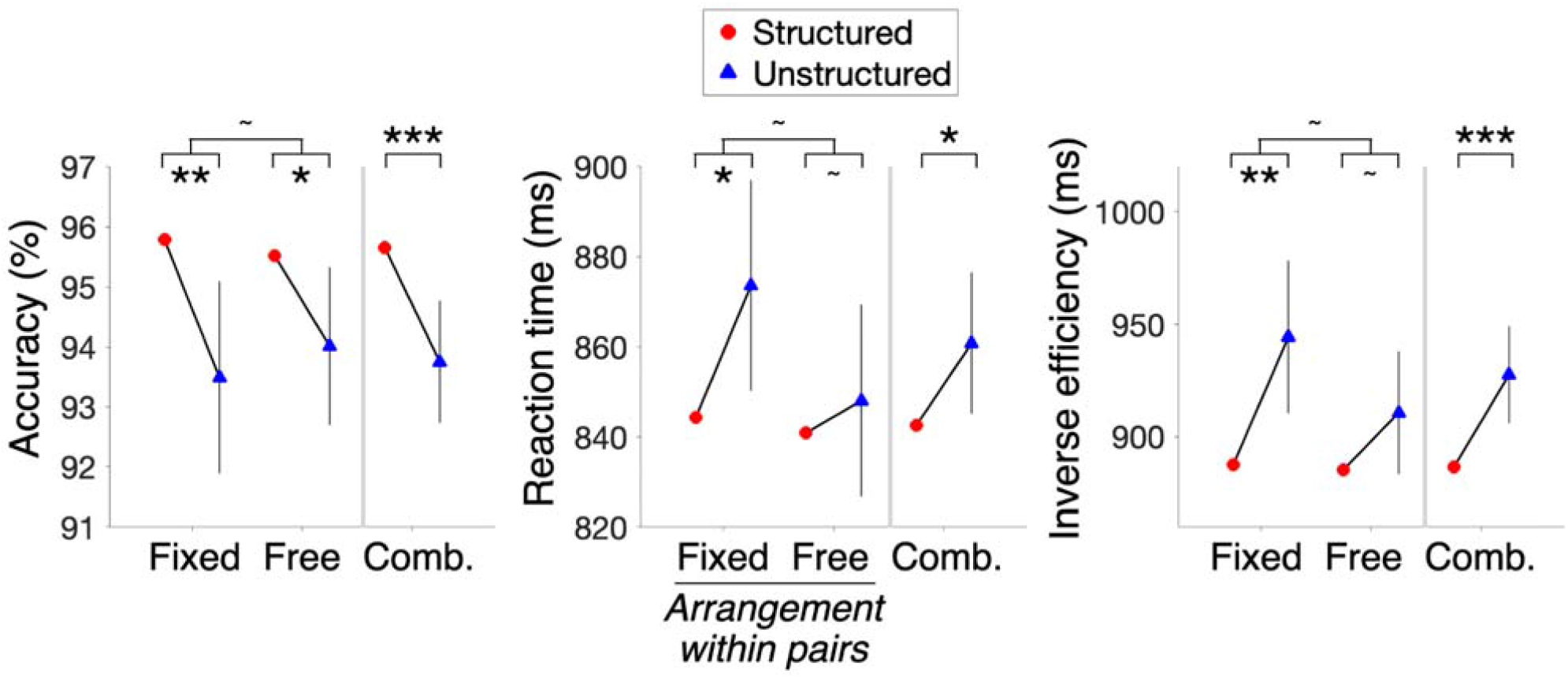
Search efficiency as a function of scene condition: Experiments 1A (Fixed) and 1B (Free). Structure-benefit (increased accuracy or decreased reaction time or decreased inverse efficiency in the structured scenes) was observed in both the experiments with fixed or free arrangements of the co-occurring shapes within their pairs. As no differences were observed between the experiments in either of the measures, the data from the two experiments were combined (‘Comb’) to accumulate the evidence for the structure-benefit. Error bars indicate 95% confidence intervals (CI) for the structure-benefit on each measure, for each experiment. Because the error bars indicate the 95% CI of the difference (structured vs unstructured), it is only shown for one of the two conditions. The asterisks indicate p-values for the t-tests for the corresponding comparisons (*p<0.05, **p<0.01, ***p<0.001, ^∼^p>0.05).

Figure 2 shows the results for Experiments 1A (Fixed arrangement) and 1B (Free arrangement). The structure-benefit did not differ across the arrangements within pairs in IES (two-sample t-test: t_78_ = 1.38, p = 0.17, d = 0.21, BF_01_ = 1.7), accuracy (t_78_ = 0.71, p = 0.48, d = 0.11, BF_01_ = 3.3) or reaction time (t_78_ = 1.36, p = 0.18, d = 0.22, BF_01_ = 1.8). Pooling across the arrangement types (denoted as ‘Comb.’ in Figure 2), there was a highly reliable structure-benefit in IES (one-sample t-test: t_79_ = 3.7, p < 0.001, d = 0.41), which was also reflected in accuracy (t_79_ = 3.6, p < 0.001, d = 0.4) and reaction time (t_79_ = 2.3, p = 0.03, d = 0.26). Thus, these experiments provided initial evidence that participants searched for targets more efficiently in the context of structured distractor arrays than unstructured distractor arrays, irrespective of the arrangement of pairs in the structured scenes.

Next, we conducted a large-sample experiment (N=400) for each of the two arrangement types with two goals in mind: First, to ensure that the structure-benefit observed in the Experiment 1 was robust (i.e., replicable in a large sample), and second, to measure participants’ familiarity for which shapes had co-occurred during the search task. Here we used one-sided t-tests to test for the existence of structure-benefits, based on the direction of the effect in Experiments 1A and 1B.

Figure 3 shows the results for Experiments 2A (Fixed arrangement) and 2B (Free arrangement). Similar to Experiment 1, the structure-benefit did not differ across arrangement type within pairs in the IES (two-sample t-test: t_798_ = 0.26, p = 0.8, d = 0.01, BF_01_ = 12.2), accuracy (t_798_ = 1.42, p = 0.16, d = 0.07, BF_01_ = 4.7) or reaction time (t_798_ = 1.18, p = 0.24, d = 0.06, BF_01_ = 6.4). Pooling across arrangement type, there was evidence for structure-benefit in the IES (one-sample, one-sided, t-test: t_799_ = 2.8, p = 0.003, d = 0.1), which was reflected both in accuracy (t_799_ = 2.5, p = 0.006, d = 0.09) and reaction time (t_799_ = 1.8, p = 0.04, d = 0.06). Thus, we found additional, confirmatory, evidence that after a period of exposure to distractor co-occurrence in the search displays, participants performed more efficient search in the structured scenes than the unstructured scenes. Notably, the benefit of distractor co-occurrence was evident irrespective of whether the co-occurring shapes in the structured scenes occurred in fixed or free arrangements within their pairs.

**Figure 3:**
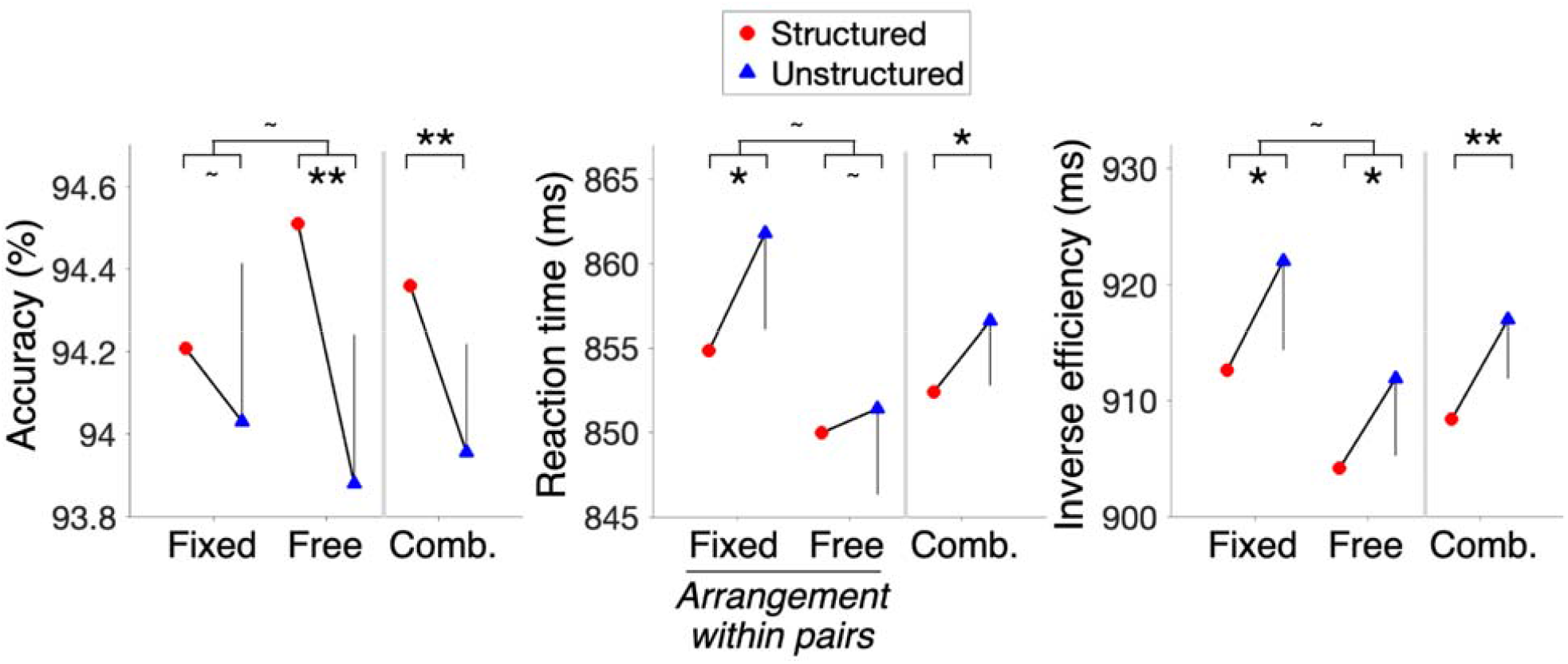
Search efficiency as a function of scene condition: Large-sample experiments 2A (Fixed) and 2B (Free). A structure-benefit (increased accuracy or decreased reaction time or decreased inverse efficiency in the structured scenes) was present for both the fixed and free pair arrangements of co-occurring shapes, replicating the effects of Experiment 1. As no differences were observed between the experiments in any measure, data from the two experiments were combined (‘Comb’) to accumulate the evidence for the structure-benefit. Error bars indicate 95% confidence intervals for the structure-benefit on each measure (corresponding to a one-sided t-test), for each experiment. Because the error bars indicate the 95% CI of the difference (structured vs unstructured), it is only shown for one of the two conditions. The asterisks indicate p-values for the t-tests for the corresponding comparisons (*p< 0.05, **p<0.01, ***p< 0.001, ^∼^p>0.05).

### Familiarity of distractor structure and its relationship with structure-benefits in search

Could participants reliably guess which pairs co-occurred during the visual search task, as has previously been reported in experiments where the pairs were passively viewed (Fiser & Aslin, 2001)? To assess whether this was the case, Experiments 2A and 2B included a 2AFC pair familiarity judgment task immediately after the main visual search task (Figure 1D). We defined *familiarity score* as the proportion of responses where the pairs corresponding to the shapes from the structured scenes were selected as more familiar than the pairs corresponding to the shapes from the unstructured scenes.

Familiarity scores for the main comparisons did not differ between Experiments 2A and 2B (two-sample t-test: t_566_ = 0.9, p = 0.4, d = 0.08, BF_01_ = 7.2). The familiarity scores did not differ significantly from 0.5 in either experiment (Experiment 2A: t_367_ = 0.85, p = 0.4, d = 0.04, BF_01_ = 11.9; Experiment 2B: t_199_ = 1.7, p = 0.08, d = 0.12, BF_01_ = 2.9), nor when we pooled the data across the two experiments for maximal power (one-sample t-test: t_567_ = 1.7, p = 0.09, d = 0.07, BF_01_ = 5.0). Finally, the two additional familiarity scores included in Experiment 2A (see Materials and Methods) also did not differ from 0.5 (position-swapped: t_367_ = 1.2, p = 0.23, d = 0.06, BF_01_ = 8.4; partner-swapped: t_367_ = 0.86, p = 0.39, d = 0.04, BF_01_ = 11.8). These results indicate that observers could not guess which shapes co-occurred during the search task.

Although the familiarity score was at chance level at the group level, it could be the case that participants who exhibited a higher structure-benefit in the visual search task were more familiar with the distractor co-occurrences, for example because they had paid more attention to these regularities during the visual search task. To test this, we assessed the correlation between the participants’ structure-benefit reflected in IES and their familiarity score. We observed a significant negative correlation when pooling the data of Experiment 2A and 2B (r = -0.10, p = 0.01). This negative correlation was significant in Experiment 2A (N = 368; r = -0.16, p = 0.001, BF_10_ = 7.3; Figure 4A), but not in Experiment 2B (N = 200; r = 0.02, p = 0.7, BF_01_ = 10.9; Figure 4B). Thus, if anything, participants who had a stronger structure-benefit in the visual search task indicated that the structured pairs were *less* familiar than the unstructured pairs in the familiarity judgment task.

**Figure 4:**
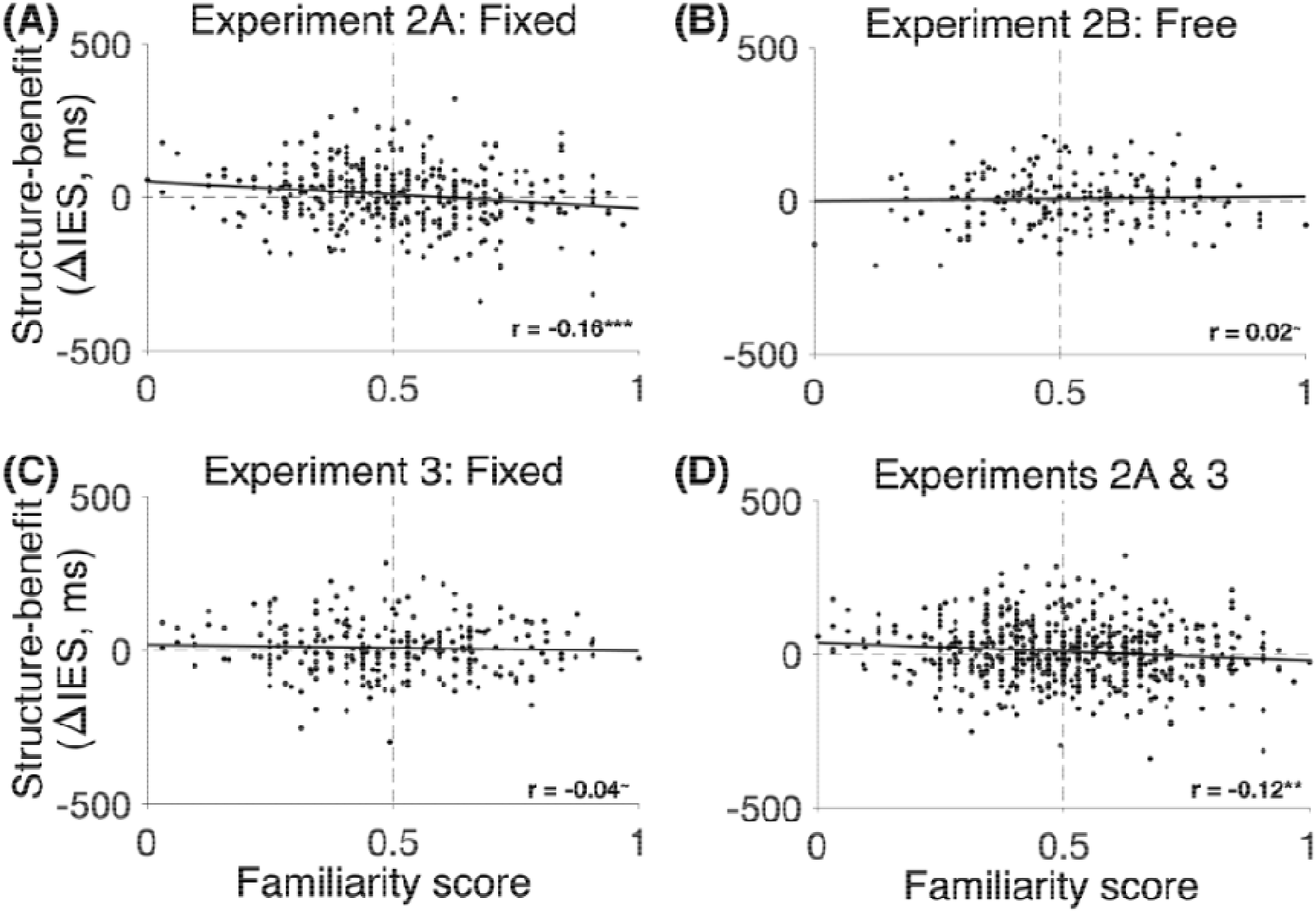
The relationship between the structure-benefit and 2AFC familiarity judgments about the co-occurring distractors. (A) In Experiment 2A, with the fixed arrangement of co-occurring distractors within their pairs, the structure-benefit (in the inverse efficiency score, IES) was negatively correlated with the familiarity scores. (B) In Experiment 2B, with the free arrangement of co-occurring distractors within their pairs, no such correlation was observed. (C) Experiment 3 did not replicate the negative correlation found in Experiment 2A. (D) Pooling across the two experiments, the structure-benefit was negatively correlated with the familiarity scores. The asterisks indicate p-values (*p< 0.05, **p<0.01, ***p< 0.001, ^∼^p>0.05).

To replicate the negative correlation of Experiment 2A, we ran a pre-registered replication of Experiment 2A (Experiment 3; N = 260).

Pre-registered analyses: The familiarity scores did not differ across comparisons (main, position-swapped, partner-swapped; F_2,518_ = 0.3, p = 0.77, BF_01_ = 34.4). Next, we created two groups of participants based on the average familiarity score: those who indicated, on average, that the pairs of objects from the structured scenes were more familiar (i.e., familiarity score > 0.5), and those who indicated the opposite (familiarity score < 0.5). Based on the results of Experiment 2A, we had pre-registered the hypothesis that the IES structure benefit would be greater for the group of participants who reported that the pairs of objects from the structured scenes were less familiar. This hypothesis was not supported by the data (one-sided t-test; t_250_= 0.45, p = 0.65, d = 0.06; BF_10_ = 0.15).

Additional analyses: As in previous experiments, the test runs of Experiment 3 demonstrated a structure-benefit in IES (one-sample, one-sided t-test: t_259_ = 1.7, p = 0.04, d = 0.11). Mirroring the findings of Experiment 2, the familiarity scores did not differ from 0.5 (one-sample t-test, main familiarity score: t_259_ = 0.3, p = 0.79, d = 0.02, BF_01_ = 13.9; position-swapped: t_259_ = 0.59, p = 0.56, d = 0.04, BF_01_ = 12.2; partner-swapped: t_259_ = 0.14, p = 0.89, d = 0.009, BF_01_ = 14.3). However, unlike Experiment 2A, the negative correlation between the main familiarity scores and the structure-benefit was not significant in this sample (r = -0.04, p = 0.44, BF_01_ = 10.5; Figure 4C).

We wondered if some difference between the responses in Experiments 2A and 3 could explain the non-replication of the negative correlation. However, there was no difference between the two experiments in either the magnitude of the structure-benefit in IES or the familiarity scores (two-sample t-tests, structure-benefit: t_626_ = 0.2, p = 0.8, d = 0.02, BF_01_ = 10.9; familiarity score: t_626_ = 0.7, p = 0.5, d = 0.06, BF_01_ = 8.4). When pooling the data across Experiments 2A and 3, the negative correlation between the structure-benefit and the familiarity scores remained significant (N = 628; r = -0.12, p = 0.003). Finally, the correlation between the structure-benefit and the familiarity scores across all available data (N = 828; Experiments 2A, 2B, 3) was also significantly negative (r = -0.09, p = 0.012).

## Discussion

Across five experiments, we found that statistical co-occurrences between distractor shapes facilitated search performance. The benefit of scene structure arose irrespective of whether the spatial arrangement of co-occurring shapes in the pairs was fixed or variable. Surprisingly, the increase in search efficiency was not accompanied by an increase in participants’ reported familiarity with the underlying statistical regularities (if anything, these effects were inversely related). These findings indicate that statistical regularities in the environment facilitate search even when these regularities are not explicitly noticed. The more efficient rejection of regular distractors may contribute to the efficiency of visual search in natural scenes, where such regularities are abundant (Kaiser et al., 2019).

How might reliable co-occurrences between distractor items give rise to a visual search benefit? Object grouping has been proposed as a complexity reduction mechanism supporting efficient search (Kaiser et al., 2014, 2019). Under this framework, shapes that consistently co-occur may be represented as a single object, similar to shapes that are grouped based on Gestalt cues (Wagemans et al., 2012). Support for this hypothesis comes from studies showing that fixed arrangements of co-occurring objects produce object-based attention effects (Lengyel et al., 2021). In our study, co-occurring distractor shapes in fixed arrangements produced more efficient search than randomly paired distractor shapes. However, a search benefit was also present (and not statistically different in magnitude) when the co-occurring shapes had *no* fixed arrangement, i.e., could vary freely in their spatial arrangement within the pair. The latter finding does not fit easily with an object grouping account, unless we assume that observers learned two objects, corresponding to the two configurations of the co-occurring shapes.

A possible alternative is that the search benefit reflected bidirectional associations between the shapes. Upon seeing one of the shapes, the representation of the associated shape may be primed, facilitating its recognition and subsequent rejection as a distractor when presented nearby. Such inter-object priming effects could operate weakly but in parallel across multiple distractor locations. The learning of arbitrary associations has been linked to the hippocampus (Davachi, 2006; Eichenbaum et al., 2007), which can modulate processing in visual cortex regions (Eichenbaum et al., 2007). Accordingly, the effects revealed here may reflect facilitated visual processing of co-occurring shapes due to hippocampus-mediated predictions (Kok & Turk-Browne, 2018). This appears to be a separate mechanism from that observed in previous studies investigating the effects of real-world positional regularities, based on long-term functional and semantic associations between objects (e.g., lamp above table). There, effects of object co-occurrences were specific to familiar spatial configurations (Kaiser et al., 2014; Quek & Peelen, 2020), and may be mediated by representational changes in visual cortex (Kaiser & Peelen, 2018) rather than hippocampus-mediated associations.

The current findings contribute to the statistical learning literature (Fiser et al., 2010) by showing that statistical regularities can be learned when these occur between shapes that have to be ignored. Unlike studies where participants passively viewed shape combinations (Fiser & Aslin, 2001, 2005), here participants could not discriminate between familiar and unfamiliar pairs post-experiment, even though the familiar pairs had been viewed 14 times more often than the unfamiliar pairs across 9 runs. This is in line with prior work that showed that such co-occurrences between items are not indicated as familiar post-experiment when the co-occurrences are task-irrelevant (Turk-Browne et al., 2005). Interestingly, if anything, the structure benefit observed in visual search performance in our study was inversely related to participants’ familiarity of the shapes. A similar negative relationship between awareness of statistical regularities and the behavioral benefit of these regularities was recently observed in a contextual cueing study, where the regularities concerned target-distractor relations (Spaak & de Lange, 2020). This suggests that statistical regularities can be learned implicitly (Turk-Browne et al., 2010). However, it is possible that familiarity would increase if the shapes were presented in the context of the original search displays. More generally, it is hard to exclude the possibility that the absence of a familiarity effect reflected the relative insensitivity of this measure (Meyen et al., 2022). We therefore interpret the dissociation between implicit and explicit measures of statistical learning with caution.

The negative correlation between structure-benefit and familiarity score suggests that participants who more effectively ignored the regular distractor pairs (thereby showing a greater structure benefit) later judged these pairs to be relatively unfamiliar. This finding may reflect the effect of inhibitory attention mechanisms, which have previously been found to suppress the visual representation of ignored objects (Seidl et al., 2012) and impair subsequent judgments on these objects (Tipper, 2001). Similarly, here, inhibiting the representations of regular distractor pairs during visual search may have resulted in these object pairs looking relatively unfamiliar during the explicit familiarity task. It should be noted, however, that the negative correlation between structure-benefit and familiarity score was not reliably observed in the pre-registered replication experiment, such that future studies are needed to confirm this account.

In summary, we find that regularities amongst distractors in the environment can be used to reduce the complexity of a scene, facilitating search for an unrelated target. Together with the encoding of regularities between distractors and targets (e.g., contextual cueing), this may help to explain the efficiency of naturalistic visual search.

## Acknowledgements

This project has received funding from the European Research Council (ERC) under the European Union’s Horizon 2020 research and innovation programme (grant agreement No. 725970).

